# Strain softening and hysteresis arising from 3D multicellular dynamics during long-term large deformation

**DOI:** 10.1101/2024.10.31.621433

**Authors:** Ken-ichi Tsubota, Shota Horikoshi, Tetsuya Hiraiwa, Satoru Okuda

**Author notes:** To whom correspondence should be addressed: Ken-ichi Tsubota (email address) and Satoru Okuda (email address).

## Abstract

Living tissues exhibit complex mechanical properties, including viscoelastic and elastoplastic responses, that are crucial for regulating cell behaviors and tissue deformations. Despite their significance, the intricate properties of three-dimensional (3D) multicellular tissues are not well understood and are inadequately implemented in biomaterial engineering. To address this gap, we developed a numerical method to analyze the dynamic properties of multicellular tissues using a 3D vertex model framework. By focusing on 3D tissues composed of confluent homogeneous cells, we characterized their properties in response to various deformation magnitudes and time scales. Stress relaxation tests revealed that large deformations initially induced relaxation in the shapes of individual cells. This process is amplified by subsequent transient cell rearrangements, homogenizing cell shapes and leading to tissue fluidization. Additionally, dynamic viscoelastic analyses showed that tissues exhibited strain softening and hysteresis during large deformations. Interestingly, this strain softening originates from multicellular structures independent of cell rearrangement, while hysteresis arises from cell rearrangement. Moreover, tissues exhibit elastoplastic responses over the long term, which are well represented by the Ramberg–Osgood model. These findings highlight the characteristic properties of multicellular tissues emerging from their structures and rearrangements, especially during long-term large deformations. The developed method offers a new approach to uncover the dynamic nature of 3D tissue mechanics and could serve as a technical foundation for exploring tissue mechanics and advancing biomaterial engineering.

## 1. Introduction

Three-dimensional (3D) biological tissues, such as organs and engineered tissues, are composed of confluent cells. These tissues exhibit characteristic mechanical responses to imposed stress and strain that are crucial for various biological processes, including embryonic development, wound healing, and disease progression [1–3]. For instance, tissues exhibit hyperelasticity and viscoelasticity, which are essential for maintaining organ structures under large deformations caused by external forces [4–7]. Additionally, tissues exhibit a solid-fluid transition that is dynamically regulated to control tissue deformability during morphogenesis [8–10]. Moreover, tissues undergo an elastoplastic transition that is adaptively regulated depending on the magnitude and duration of an applied deformation to ensure unidirectional progression of morphogenesis [11]. These properties arise from hierarchical tissue structures and their interactions, ranging from molecular and cellular constituents to the entire tissue. However, our understanding of the mechanical mechanisms underlying 3D tissue properties remains limited due to a lack of methods to analyze these properties based on the 3D hierarchical structures and dynamics of multicellular tissues. Therefore, development of methods to analyze the multicellular behaviors within 3D confluent cell tissues and understanding of their fundamental properties are essential to advance the biological understanding and biomaterial engineering of tissues [12,13].

Advances in measuring tissue mechanics have significantly enhanced our understanding of the mechanisms that regulate their properties. A pioneering study measured the stiffness of whole *Xenopus laevis* embryos using micro-aspiration, which revealed that embryo stiffness varies during gastrulation [14]. Rheological properties of multicellular aggregates have been assessed using rheometers, revealing nonlinear rheology [15,16]. Atomic force microscopy has also been used to measure tissue stiffness, showing spatiotemporal changes during mouse brain and retina development [17,18]. Additionally, magnetic oil droplets and thermally responsive gels injected into tissues have been used to successfully measure viscoelastic properties within embryonic tissues [5,19,20]. These measurements have highlighted the diverse mechanical responses of 3D multicellular tissues, including viscoelasticity and hysteresis. Recent studies have further revealed that cells exhibit both solid-like and fluid-like characteristics, along with transitions between these states [9,10,21,22]. These properties arise from the confluent cell structure within multicellular tissue and are profoundly influenced by cell rearrangements [21,23,24], which are regulated by junctional adhesion molecules [25,26]. Meanwhile, tissues also exhibit plastic responses that are directly regulated by actin polymerization [11,18]. Thus, recent reports have identified several key cellular behaviors crucial for determining tissue properties. However, there remains a significant gap in our knowledge regarding how these cell behaviors integrate to determine properties of the entire tissue.

To explore the mechanical mechanisms regulating tissue properties, we herein developed a new mathematical method for mechanical tests using a 3D vertex model framework [27,28]. This method addresses three major requirements. First, the model needs to have a single-cell resolution to capture the hierarchical structure and dynamics of tissues from the subcellular to the tissue level, such as actomyosin contractility, cadherin-induced cell-to-cell adhesion, and 3D cell rearrangement. Cell-based models, such as particle models [29–32] and vertex models [33,34], could be applied to analyze such tissue dynamics with single-cell resolution. Second, the description of the tissue structure needs to be in 3D space. Although mathematical studies of multicellular dynamics have been extensively conducted for 2D tissues and cell monolayers [35–37], fundamental differences between 2D and 3D models may lead to distinct properties. For instance, at the continuum level, Poisson effects and anisotropy in tissue properties can be significant [38,39]. At the cellular level, differences exist in the topological paths of cell rearrangements [40], making it essential to understand their role in tissue properties. To analyze such 3D tissues with single-cell resolution, 3D models such as 3D vertex- and 3D Voronoi-based models [27,41–43], which describe the 3D multicellular structure and rearrangement [27,28,41], could be applied. Third, the model needs to be applied to general tests of mechanical properties, such as stress relaxation tests and dynamic viscoelastic analysis [44]. Previously, we reported tensile and compressive tests of epithelial tubes using a 3D vertex model framework, revealing the effects of actomyosin distribution on tissue responses [45,46]. However, although several methods exist for studying 3D multicellular dynamics, the rheological properties of confluent 3D tissues remain largely unknown, and there is still a significant lack of methods that can be specifically applied to confluent 3D tissues. Therefore, engineering of 3D multicellular tissues requires methods that can accurately describe and predict their properties based on single-cell 3D behaviors. Our new framework aims to fill this gap, providing a comprehensive approach to analysis of the mechanical properties of 3D confluent cell tissues.

To demonstrate mechanical tests of 3D confluent cell tissues using a 3D vertex model, we conducted stress relaxation tests and dynamic viscoelastic analyses. While multicellular tissues maintain their 3D structures in response to external forces, they undergo large deformations over long timescales, such as during embryogenesis. Although cell rearrangement is important for long-term large deformations [28,47], the mechanisms contributing to tissue properties remain unclear. To the best of our knowledge, there are no reports of theoretical analyses of 3D tissue properties during long-term large deformations. Here, we first performed stress relaxation tests to clarify the stress response to deformation arising from cell rearrangement by comparing results with and without cell rearrangement. We analyzed the mechanisms underlying this response by quantifying the frequency of cell rearrangements and the sphericity of individual cells during the relaxation process. Next, we conducted dynamic viscoelastic analysis to reveal strain softening and hysteresis of the stress response in relation to tissue fluidization. Quantitative comparisons of the stress-strain relationship, frequency dependence, and individual cell dynamics revealed different mechanisms underlying strain softening and hysteresis. Finally, we applied the Ramberg–Osgood model [48] to quantitatively characterize the long-term response, including strain softening and hysteresis. Our new method sheds light on the mechanical properties of 3D confluent cell tissues that emerge from their hierarchical structure and dynamics and serves as a general tool for the broader field of tissue mechanics and biomaterial engineering.

## 2. Methods

### 2.1 Three-dimensional vertex model framework

To develop a method to measure 3D tissue properties, we introduced a 3D vertex model to describe multicellular dynamics in 3D tissues [28]. In this model, individual cells are represented as polyhedrons with vertices shared by neighboring cells (Fig. 1**a**). The shape and configuration of the cells are determined by the positions of the vertices comprising the cellular polyhedrons and the topological network among these vertices.

**Fig. 1.**
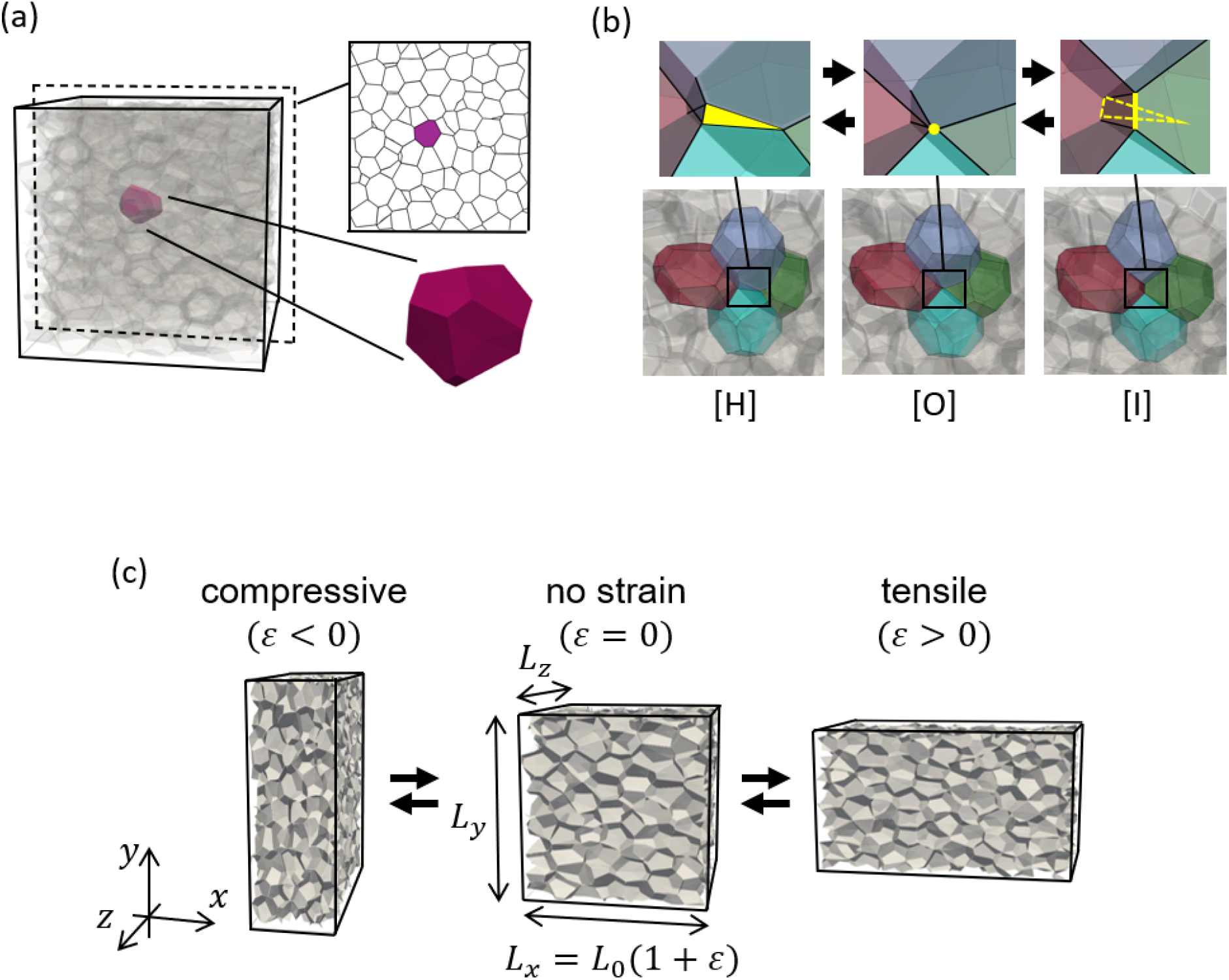
Three-dimensional vertex model of multicellular tissue. (a) Cells consist of vertices, edges, and faces, which are shared with neighboring cells. (b) Cell-to-cell rearrangement is shown. The [H]-to-[I] operation replaces the triangular surface in [H] with the edge in [I] via the point in [O], as indicated by yellow-colored parts. The [I]-to-[H] operation occurs vice versa. (c) The shapes of a tissue in the mechanical test are shown. The entire tissue is represented as a rectangle, and the lengths in each direction are *L*_*x*_, *L*_*y*_, and *L*_z_ (middle). Compressive (left) or tensile (right) strain is applied in the *x* direction in the mechanical test, where the length in the *x* direction is described as *L*_*x*_ = *L*_0_(1 + *ε*)with the axial strain *ε*.

Cell movements are expressed by changes in vertex locations. The time evolution of the *i-*th vertex location, represented by ***r***_*i*_, is given by

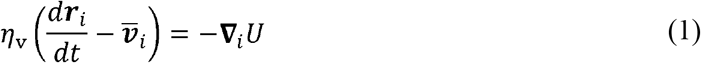

The left side of Eq. (1) represents the friction force acting on the *i-*th vertex [37]. Here, we assume that friction forces are exerted between adjacent cells, *η*_v_ is the friction coefficient, defined as the summation of the friction coefficients from the adjacent cells sharing cell-cell boundaries: 4*η*_c_, where *η*_c_ is the friction coefficient between cells. The vector 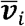 is a velocity field defined as the mean velocity of the surrounding cells: 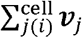, where ***v***_*j*_ is the velocity of the *j*-th cell, calculated as the mean velocity of the vertices composing the *i*-th cell. The right side of Eq. (1) represents the mechanical force acting on the *i*-th vertex, derived from the effective energy, denoted by *U*. **∇**_*i*_ denotes the gradient with respect to ***r***_*i*_.

During cell movements, as described by Eq. (1), individual edges and polygons in the network occasionally shrink, either to meet or retract from neighboring cells. Corresponding to those vertex movements, cell rearrangement is performed, defined as the dynamic reconnection of the topological network using [I]-to-[H] and [H]-to-[I] operations [49] (Fig. 1**b**). These operations between patterns [H] and [I] are executed when the related edge lengths become infinitesimally small, ensuring energy conservation during the operations [28]. The [H]-[I] operations represent changes in 3D cell configurations, differing from the well-known T1 transformation used in 2D models.

### 2.2 Energy function, system box, and stress tensor

While focusing on a 3D tissue composed of confluent homogeneous cells, we describe cell behaviors by introducing a simple effective energy, *U*, on the right side of Eq. (1):

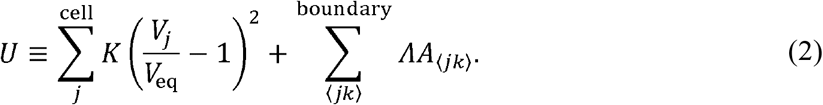

The first term represents the volume elastic energy, where *K* is an elastic constant and *V*_*j*_ is the volume of the *j*-th cell. The second term represents the boundary energy, where *Λ* is the area tension on the ⟨*jk*⟩-th boundary with area *A*_⟨*jk*⟩_. The symbol ⟨*jk*⟩ denotes the boundary polygon between the *j-*th and *k-*th cells. *Λ* represents the average surface tension exerted on the boundary polygon, corresponding to the summation force exerted on the cell-cell boundary, such as in cortical actin cytoskeletons and interfacial adhesions.

Using *V*_eq_, *K*, and *η*_c_, all parameters are nondimensionalized by units: length *l* = (*V*_eq_)^1/3^, energy *e* = 0.1 *K*, and time *τ* = 20 *η*_c_ (*V*_eq_)^2/3^/K. The common set of simulation parameters includes the equilibrium volume of cells *V*_eq_ = *l*^3^, elastic constant of volumetric energy *K* = 10 *e*, friction coefficient *η*_c_ = 0.5 *τe*/*l*^2^, and intercellular tension *Λ* = 0.2 *e*/*l*^2^.

To perform stress relaxation tests and dynamic viscoelastic analyses, we utilized a simple system of compacted cells within a box (Fig. 1**c**). The box dimensions are defined within 0 ≤ *α* ≤ *L*_*α*_ (*α* = *x, y, z*), where periodic boundary conditions are imposed at *α* = 0 and *L*_*α*_. Each cell’s volume is set to be approximately *V*_eq_, resulting in the total volume of the box being constrained to *L*_*x*_*L*_*y*_*L*_*z*_ = *N*_t_*V*_eq_, where *N*_t_ is the total number of cells in the box. For the stress relaxation tests and dynamic viscoelastic analyses, axial strain *ε* in the *x*-direction was applied (Fig. 1**c**). The length of the box in the *x* direction, *L*_*x*_, is given by *L*_*x*_ = (1 + *ε*), where *L*_0_ = (*N*_t_*V*_eq_)^1/3^ is the length of the box with no strain (*ε* = 0).

In stress relaxation tests and dynamic viscoelastic analyses, the stress response of the tissues was measured by uniaxial mechanical testing, which assumes affine deformation under the constant volume *L*_*x*_*L*_*y*_*L*_*z*_ = *N*_t_*V*_eq_. The stress tensor of the system, denoted by *σ*_*αβ*_, is calculated by

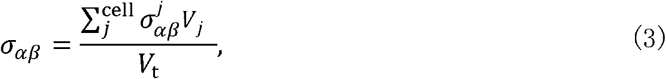

where the summation is performed over all cells. In this equation, 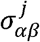 refers to the stress tensor of the *j*-th cell. We define 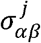 as

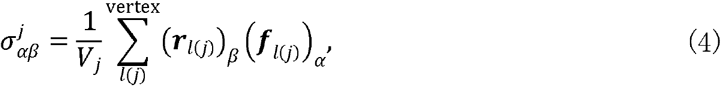

where the summation is performed over all vertices composing the *j*-th cell. Here, ()_*α*_indicates the *α*-th component of a vector. The vector ***r***_*l*(*j*)_ is the distance vector from the center of the *j*-th cell to the position of the *l*-th vertex. ***f***_*l*(*j*)_ represents the force acting on the *l*-th vertex within the *j*-th cell, given as ***f***_*l*(*j*)_ = −*dU*_*j*_/*d****r***_*l*_. The normal stress *σ*_*xx*_ of the loading direction was considered as the representative stress, and hereafter, it is described simply as *σ*.

### 2.3 Stress relaxation test

Stress relaxation tests were conducted to clarify viscoelastic properties under simple loading conditions. Axial strain *ε* in the *x* direction was applied to the tissue at time *t* = 0 and maintained until the tissue reached mechanical equilibrium (Fig. 2**a**). To analyze the effects of cell rearrangements and the magnitude of strain *ε* on tissue responses, simulation parameters were varied with, *ε*= 0.1–0.5, both with and without cell rearrangements.

**Fig. 2.**
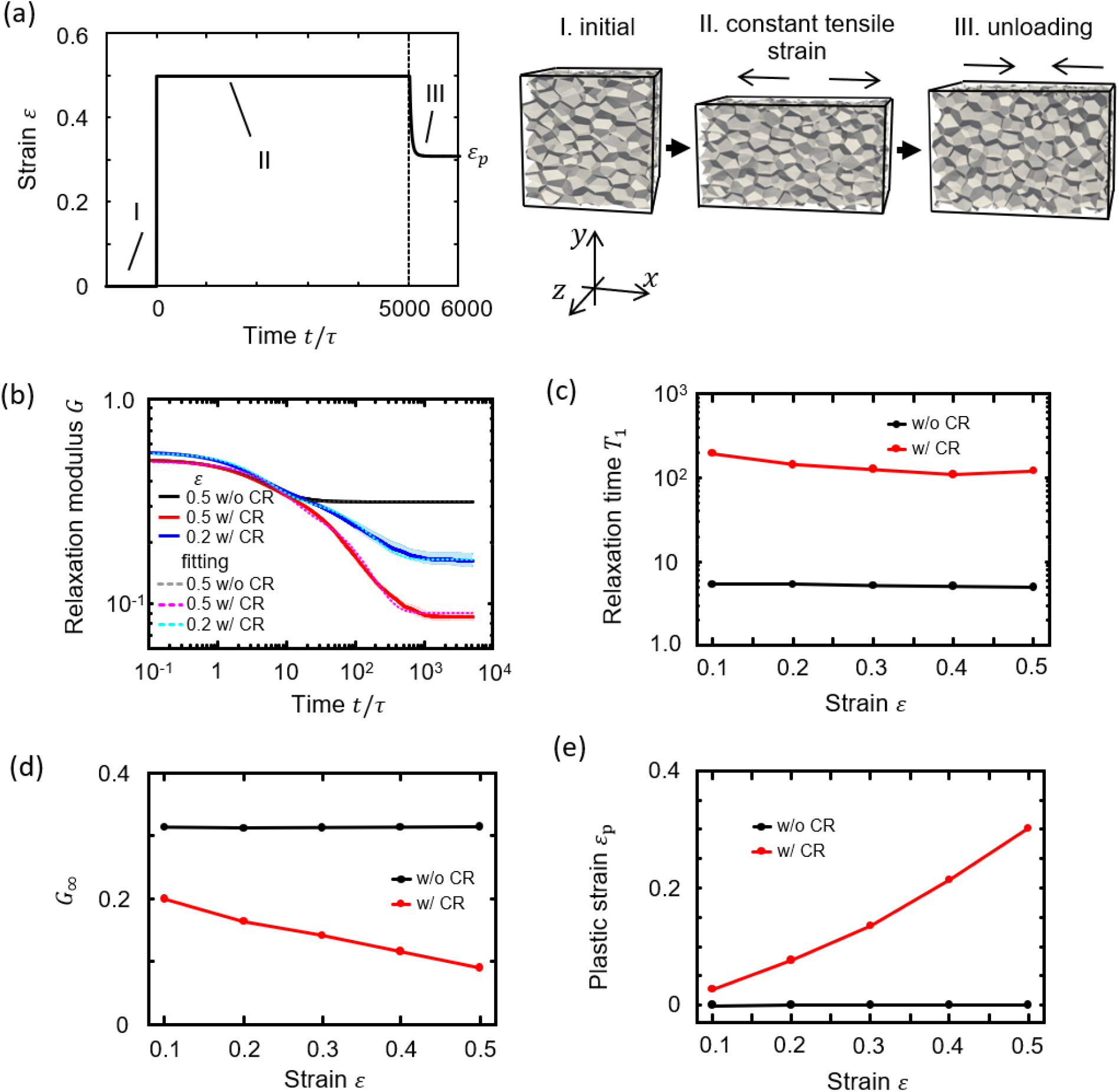
Viscoelastic and plastic properties obtained in stress relaxation tests. (a) Strain *ε* is instantaneously applied at time *t* = 0 (from state I to II) and maintained until the mechanical equilibrium is obtained (state II). The unloading test is performed after stress relaxation (state III). (b) Time-course of changes in relaxation modulus *G* (solid lines) in cases without cell rearrangement (w/o CR, black) and with cell rearrangement (w/ CR, red) at strain *ε* 0.5 obtained in the stress relaxation tests. Data for the case with cell rearrangement at strain, *ε =* 0.2 (blue) are also illustrated. The band widths of the light colors indicate the standard deviation. Fitting of *G*(*t*) by the generalized Maxwell model (Eq. (5)) is illustrated by dashed lines. (c) Relaxation time *T*_1_ and (d) elastic modulus *G*_*∞*_ as a function of strain *ε* in cases without cell rearrangement (w/o CR, black circles) and with cell rearrangement (w/ CR, red circles). (e) Plastic strain *ε*_P_ obtained in the unloading test as a function of strain *ε*.

The stress response was analyzed by calculating the relaxation modulus *G*, defined as *σ*/*ε*, cellular deformation according to the sphericity of each cell, represented by *S*, and cell rearrangement according to the number of cells adjacent to each cell, represented by *N*_AC_. Additionally, to analyze the relaxation process, *G* was fitted by the generalized Maxwell model [50], which is a parallel association of a linear spring and *N*_Maxwell_ sets of a linear spring and damper defined as

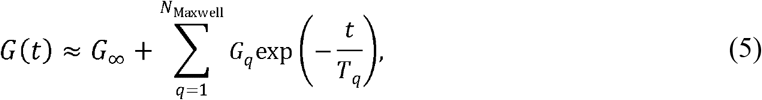

where *G*_∞_ indicates the relaxation modulus in mechanical equilibrium, *G*_*q*_ and *T*_*q*_ and *T*_*q*_ (*T*_*q*_ ≥ *T*_*q*+1_) are elastic constants and relaxation times, respectively. The parameters *G*_∞_, *G*_*q*_, and *T*_*q*_ were determined by fitting the simulation results with Eq. (5).

To evaluate the plastic property of tissues, we further conducted an unloading test of the multicellular tissue after the stress relaxation test and quantified the plastic strain *ε*_p_ generated in plastic strain *ε*_p_ was defined as 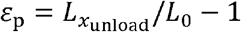, where 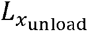 is the length *L*_*x*_ at the final the stress relaxation test. During the unloading test, we did not implement cell rearrangements. The state of the unloading test.

### 2.4 Dynamic viscoelastic analysis

For dynamic viscoelastic analyses, a cyclic tension-compression test was conducted to assess the mechanical responses of tissues to a wide amplitude and frequency range. Dynamic strain was applied as *ε* = *ε*_A_ Sin(2*πft*), where *ε*_A_ and *f* represent the strain amplitude and frequency, respectively. To analyze the effects of cell rearrangements and the magnitudes of strain *ε* and frequency *f* on tissue responses, simulation parameters were varied with strain amplitude *ε*_A_ ranging from 0.1 to 0.5 and regularized strain frequency *fτ* ranging from 2.0 × 10^−4^ to 0.1, both with and without cell rearrangements.

Dynamic loadings were repeated for five cycles after eliminating the effects of initial conditions on mechanical behaviors by using preliminary loadings for several cycles. Using the data from these five cycles, the axial stress *σ* was fitted by the Maxwell model as

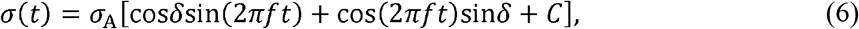

using the least squares method. In Eq. (6), *σ*_A_, *δ*, and *C* are the stress amplitude, initial phase, and constant determining time-averaged stress, respectively, as the parameters to fit. Subsequently, the storage modulus *G*′ (corresponding to elasticity), loss modulus *G*″ (viscosity), and loss tangent tan*δ* (viscosity relative to elasticity) [50] were determined using the constants *ε*_A_, *σ*_A_, and *δ* as

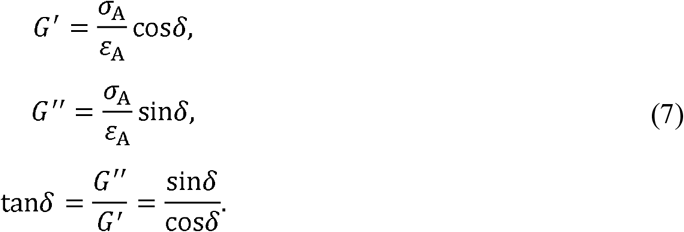

and

The moduli *G*′, *G*″, and tan*δ* were given as a function of strain frequency *f*.

To analyze the mechanical response of tissues, we fitted moduli *G*′, *G*″, and tan*δ* as a function of strain frequency *f* using the generalized Maxwell model with a linear spring and *N*_Maxwell_ sets of a linear spring and damper:

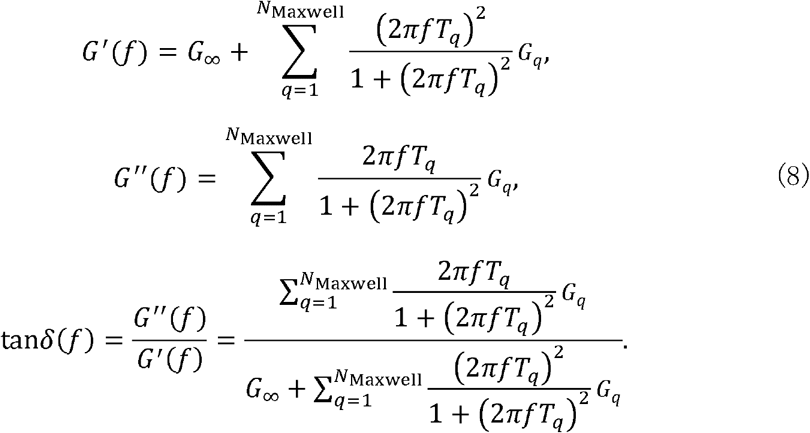

and
In Eq. (8), the elastic constants *G*_∞_ and *G*_*q*_ and relaxation time *T*_*q*_ are the parameters to fit. In addition, stress *σ*(*t*) simulated at a certain frequency *f* of dynamic strain was transformed into the frequency domain to obtain the imaginary stress amplitude *σ*_A(*k*)_ as a function of wavenumber *k* using a discrete Fourier transform. Using cyclic tension-compression data for *m* cycles for which the duration time is *m*/*f*, the Fourier transform was calculated as

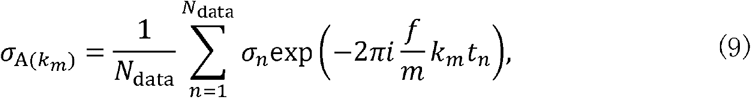

where *k*_*m*_ is the wavenumber with respect to duration time *m*/*f, σ*_*n*_ is the stress at time *t*_*n*_ of the *n*-th simulation data (*n* = 1,…, *N*_data_), and *i* is the imaginary unit. Using the condition *k* = *k*_*m*_/*m*, which is an integer, the stress amplitude σ_A(*k*)_ in the frequency domain with respect to the strain period 1/*f* was extracted from 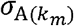.

### 2.5 Model fitting

Parameters involved in Eqs. (5), (6), and (8) were determined by numerically searching a set of parameters that minimize the sum of the squares, 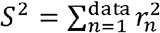, of the residuals *r*_*n*_ about the *n*-th simulation data. The squares 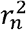 of the residual were defined for Eqs. (5), (6), and (8), respectively, as

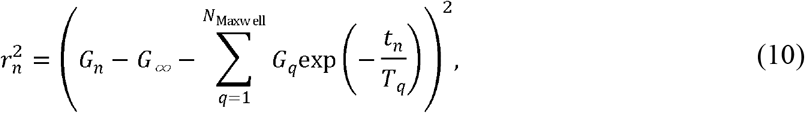

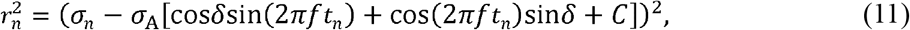

and

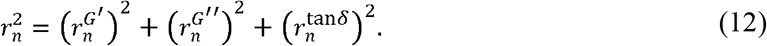

Here, 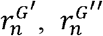 and 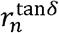 in Eq. (12) for fitting Eq. (8) are denoted as

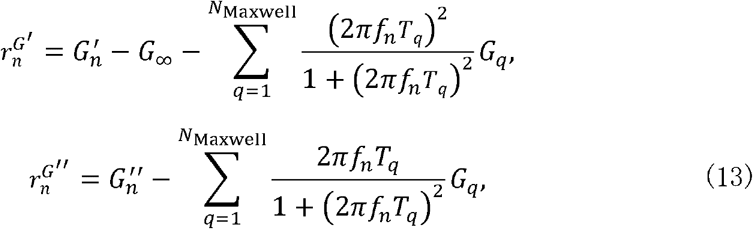

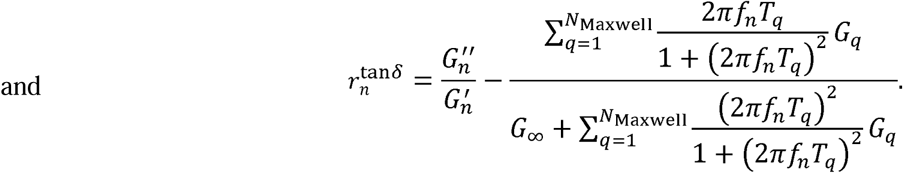

### 2.6 Numerical procedures and statistics

Vertex velocities were implicitly calculated by simultaneously solving Eq. (1) for all vertices. The time development of vertex movements was calculated by integrating Eq. (1) using the Euler method with a time step of Δ*t* (= 0.005*τ*). The [I]-to-[H] and [H]-to-[I] operations were applied at every time step of Δ*t*_r_ (= *τ*), where the threshold length for these operations was set to Δ*l* (= 0.05 *l*).

The initial condition was prepared as follows: the box was initially filled with 128 cells, and cell proliferation was simulated over two cell cycles [51]. The box size was then deformed in an affine manner to relax the normal stress along each of the *x*-, *y*-, and *z*-axes to zero. The resulting box contained a total of *N*_t_ = 536 ± 8.9 (mean ± standard deviation) cells, with slight variation depending on the test specimen caused by randomized cell cycles. For all parameter sets, we performed five individual simulations with different initial cell structures, and the mean and standard deviation of the simulation data over the five individual simulations were calculated. The mean sphericity of cells under this initial condition, denoted by *S*_0_, was approximately *S*_0_ = 0.90, closely approximating the unity of a sphere. The simulations were carried out five times for each parameter, each with different initial structures of cells, and statistical data were calculated based on these simulations.

## 3. Results

### 3.1 Stress relaxation tests unveil quantitative impacts of cell rearrangement on tissue viscoplastic responses

To investigate the mechanical mechanisms that regulate the properties of 3D confluent cell tissues, it is essential to understand the entire tissue response to the applied stress or strain based on individual cell behaviors. However, conducting mechanical tests of 3D multicellular dynamics using 2D or continuum approximations is challenging. To overcome this problem, we developed a method to conduct mechanical tests of 3D confluent cell tissues using a 3D vertex model. We chose a tissue composed of cells with uniform mechanical behaviors, including volume elasticity and cell-cell interfacial tension. To conduct mechanical tests on 3D bulk tissues, the stress of the entire tissue under periodic boundary conditions needs to be observed. We defined the stress tensor under these conditions as shown in Eq. (3), where we first calculate the stress tensor within each cell and then average it over all cells. Using this procedure, the stress tensor of the entire tissue can be calculated, even under periodic boundaries.

We performed stress relaxation tests by preparing the initial condition of a tissue under stress relaxation and then applying a constant uniaxial strain *ε* along the *x*-axis under incompressible conditions (state I and state II in Fig. 2**a**). By observing the stress relaxation process, we obtained the relaxation modulus *G* (= *σ*/*ε*) as a function of time, which decreased over time and reached a constant value (Fig. 2**b**). To clarify the effects of *ε* and cell rearrangement on the relaxation process, we compared the results under conditions with high and low *ε*, both with and without cell rearrangement. The decrement in *G* increased with *ε* and with cell rearrangement. Specifically, in cases of high *ε, G* decreased similarly in cases with and without cell rearrangement in a short timescale (*t*/*τ* < 10), but the decrement in *G* was accelerated by cell rearrangement over a long timescale (10 < *t*/*τ*). These results show that 3D confluent cell tissues exhibit stress relaxation, which can be clearly divided into two relaxation processes: a short timescale process independent of cell rearrangement and a long timescale process caused by cell rearrangement.

To evaluate these relaxation processes, we fitted using the Maxwell model (dashed lines in Fig. 2**b**), where *N*_Maxwell_ = 1and 2 (Eq. (5)) in cases without and with cell rearrangement, respectively, and obtained the longest relaxation time *T*_1_ and relaxation modulus under equilibrium *G*_∞_. The difference between cases in the presence and absence of cell rearrangement (more than 10 times) was much greater than that for various *ε* values within each case (maximum two times) (Fig. 2**c**). Moreover, while *G*_∞_ remained constant with respect to *ε* in cases without cell rearrangement, *G*_∞_ decreased with increasing *ε* in cases with cell rearrangement (Fig. 2**d**).

To understand the mechanism causing the difference between cases with and without cell rearrangement, we conducted an unloading test after the stress relaxation test, where *ε* was relaxed to reach *σ* = 0 (state III in Fig. 2**a**). The obtained plastic strain *ε*_p_ was zero in cases without cell rearrangement, independent of *ε*. However, *ε*_p_ increased with *ε* with cell rearrangement (Fig. 2**e**). These results show that cell rearrangement allows plastic deformation, occurring over a certain long-term scale, with the amount of plastic deformation increasing with *ε*.

Recent studies suggest the impact of cell rearrangements on tissue properties [5,21,23,24]. However, while it is known that cell rearrangement occurs during the stress relaxation process, its quantitative contributions remain largely unknown. Our method clarified the viscoplastic property of 3D confluent cell tissues and quantified the effects of cell rearrangement on this property. Notably, we found that *T*_1_ increased by 20 times with cell rearrangement over a certain period, independent of *ε* (Fig. 2**c**), *G*_∞_ decreased to one-third due to cell rearrangement (Fig. 2**d**), and the ratio of plastic deformation to the applied deformation exceeded 50% (e.g., *ε*_p_, ∼0.3 when *ε* = 0.5) (Fig. 2**e**). A recent study demonstrated that plastic deformation occurs when the applied maximal deformation is approximately 50% or more of the average cell size [5], corresponding to *ε* ∼0.5. Therefore, our simulation results align with this recent study. These findings underscore the quantitative impact of cell rearrangement on the stress relaxation of multicellular tissues.

## 3.2 Cell shape recovery, enhanced by cell rearrangement, drives tissue stress relaxation

To address the mechanisms that determine stress relaxation in 3D confluent cell tissues, we focused on cell shapes and rearrangements during the relaxation process. Cell behaviors are described by the simple formulation in Eq. (2), which induces shape resilience in each cell as a result of the force balance between volume elasticity and surface tension. To assess cell shapes, we calculated the mean value of cell sphericity *S* (Fig. 3**a**), its distribution across cells (Fig. 3**b**), and the mean sphericity after reaching a constant state during relaxation, *S*_∞_ (Fig. 3**c**).

**Fig. 3.**
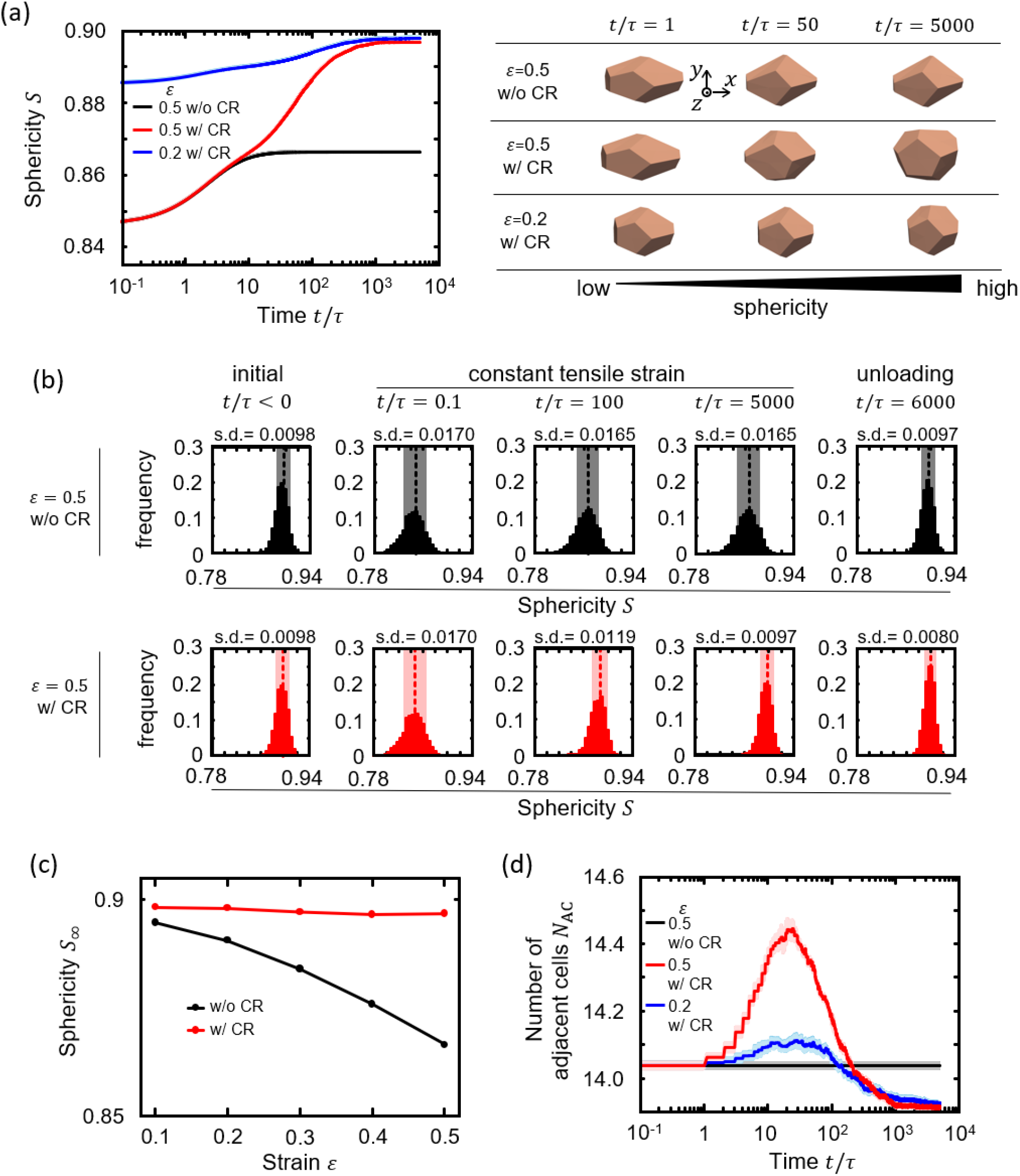
Cellular structures obtained in stress relaxation tests. (a) Time-course of changes in sphericity *S* in the cases of *ε* = 0.5 without cell rearrangement (black line) and with cell rearrangement (red line) and of *ε* = 0.2 with cell rearrangement (blue line). The standard deviation for all cells in the five individual simulations is less than 0.1% of the *S* value and thus not visible in the panel. The representative cellular shapes are also illustrated for the early (*t*/*τ* = 1), middle (*t*/*τ* = 50), and final (*t*/*τ* = 5000) stages of the relaxation processes. (b) Distribution of sphericity *S* inside the multicellular tissue in the case of *ε* = 0.5 without cell rearrangement (upper row) and with cell rearrangement (lower row) at the initial shape (*t*/*τ* < 0) under constant tensile strain (*t*/*τ* = 0.1,100,and 5000) and at unloading (*t*/*τ* = 6000). The dashed lines in each panel indicate the mean values of *S*, and the band widths of light colors indicate the standard deviation. (c) Sphericity at the mechanical equilibrium, *S*_∞_, as a function of strain *ε*. The cases without cell rearrangement (w/o CR, black circles) and with cell rearrangement (w/ CR, red circles) are shown. (d) Time-course of changes in the number of adjacent cells *N*_AC_. The same line colors as in Fig. 3**a** are used to indicate *ε* and conditions with and without cell rearrangement. The band widths of the light colors indicate the standard deviation.

Initially, the mean cell sphericity *S*, which was *S*_0_ (= 0.90) in the absence of strain, decreased as the initial strain *ε* increased (Fig. 3**a**). After the initial strain was applied, *S* increased over time, both in cases with and without cell rearrangement, eventually reaching a steady-state value, *S*_∞_. In the case with cell rearrangement, *S* reached a higher value, closer to *S*_0_, than that in the case without cell rearrangement (Fig. 3**a**). The final sphericity *S*_∞_ was approximately equal to *S*_0_ regardless of *ε* when cell rearrangement occurred, whereas it decreased with increasing *ε* in the absence of rearrangement (Fig. 3**c**). These results suggest that the initial strain applied to tissue is primarily restored in each cell and that relaxation is drastically facilitated by cell rearrangement.

Cell sphericity varied across individual cells, appearing to follow a normal distribution centered around the mean value (Fig. 3**b**). The variance in cell sphericity increased due to the initial strain (from *t*/*τ* = 0 to 0.1) in both conditions, with and without cell rearrangement. During relaxation (from *t*/*τ* = 0.1 to 5,000), the variance remained high in the absence of cell rearrangement, while it gradually narrowed in the case with cell rearrangement. Upon unloading, the variance narrowed in both conditions, although the distribution in the case with cell rearrangement was more concentrated than that in the case without rearrangement. These findings suggest that cell rearrangement plays a crucial role in homogenizing the shapes of individual cells.

To clarify the relationship between cell shape and rearrangement, we calculated the number of cells adjacent to each cell, *N*_AC_. *N*_AC_ started to change at approximately *t*/*τ* = 2 and converged to a constant at *t*/*τ* = 2,000 (Fig. 3**c**). The change in *N*_AC_ began earlier than the deviation in cell sphericity between cases with and without cell rearrangement, which started at *t*/*τ* = 10 (Fig. 3**a**,**d**). However, the convergence timing of *N*_AC_ at *t*/*τ* = 2,000 was consistent with that of sphericity. These observations demonstrate that cell shape recovery, enhanced by cell rearrangement, drives tissue stress relaxation, leading to viscoplastic tissue behaviors.

Our new method enabled quantitative analyses of the mechanical properties of 3D confluent cell tissues. It was particularly effective in analyzing the time dependency of tissue responses and understanding tissue properties based on cell behaviors. We will explore the time dependency of tissue responses and their underlying mechanisms in the following sections.

### 3.3 Dynamic viscoelastic analyses reveal apparent tissue fluidization due to strain softening during long-term large deformation

To investigate dynamic properties of 3D confluent cell tissues, we conducted a dynamic viscoelastic analysis by applying cyclic uniaxial tension-compression to the tissue with dynamic strain *ε* = *ε*_A_ Sin(2*πft*) along the *x*-axis (Fig. 4**a**). We obtained stress *σ* as a function of time, which exhibited steady oscillation in response to *ε* (Fig. 4**b**). In the case without cell rearrangement, *σ* followed strain *ε* without a phase shift, maintaining a sine curve. However, *σ* became asymmetric between positive and negative values due to the difference between the tensile and compressive stress in uniaxial deformation. In contrast, in the case with cell rearrangement and large *ε*_A_ (= 0.5), *σ* exhibited a phase advance and deviation from a sine curve, indicating hysteresis and nonlinearity of the response. The phase advance increased with *ε*_A_, as is illustrated by the difference in *σ* between *ε*_A_ = 0.2 and *ε*_A_ = 0.5, reflecting the viscosity enhancement driven by the plasticity due to cell rearrangement as shown in Fig. 2**e**.

**Fig. 4.**
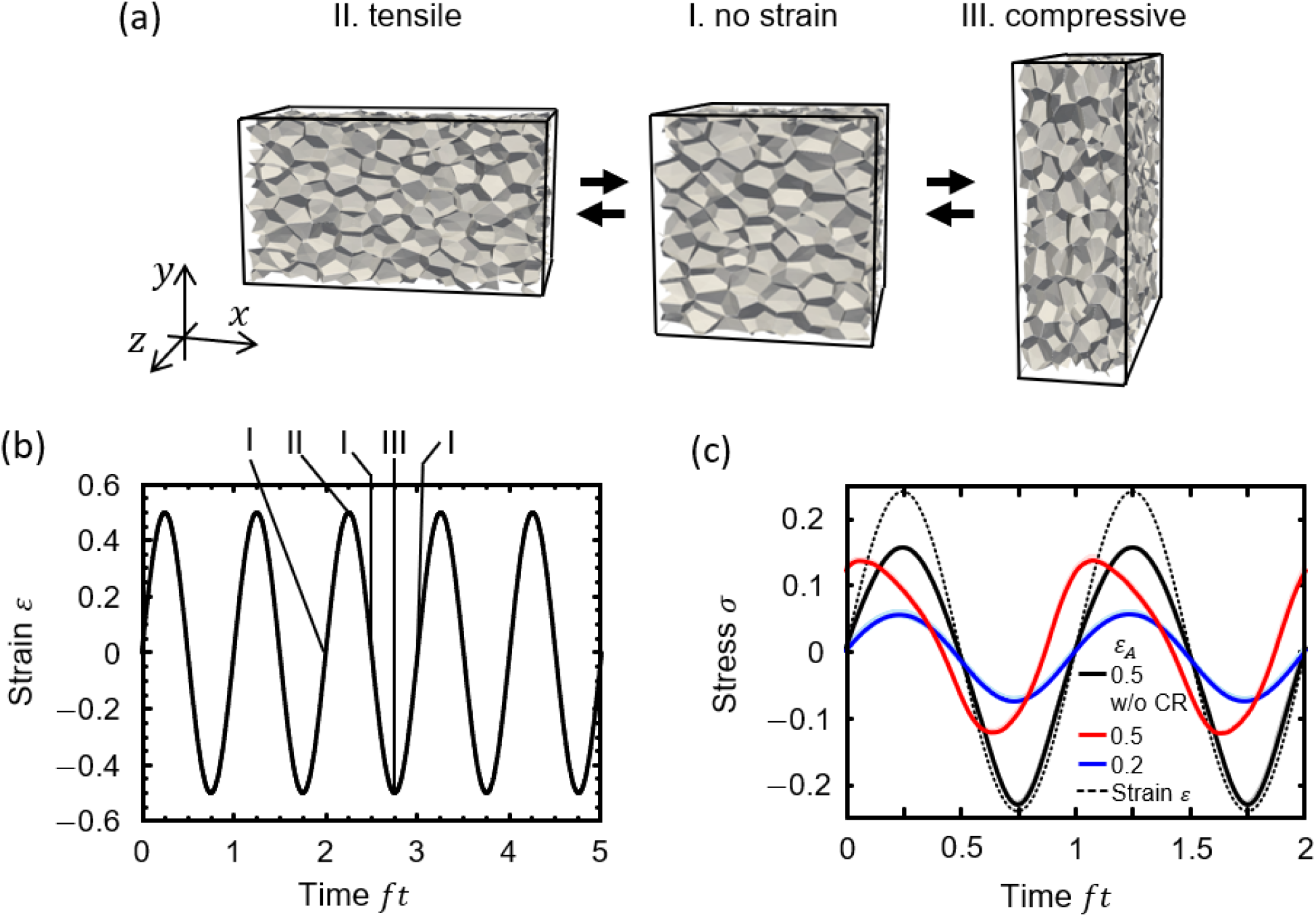
Simulation of dynamic mechanical analysis (*fτ* = 0.001). (a) Cellular shapes in cyclic strain paths consisting of no strain (I)–tensile strain (II)–no strain (I)–compressive strain (III). (b) Applied strain *ε* as a function of time *ft*. The deformation states I, II, and III correspond to the cellular shapes shown in Fig. 4**a**. (c) Time-course of changes in stress *σ* for cases without cell rearrangement (black line) and without cell rearrangement (w/o CR, red line) at a strain amplitude *ε*_A_ = 0.5 and with cell rearrangement at *ε*_A_ = 0.2 (blue line). Strain is also illustrated (dotted line) for comparison.

To quantify the time dependence of viscoelastic properties, we calculated the storage modulus *G*′, loss modulus *G*″, and loss tangent tan*β* as a function of strain frequency *f* both in cases with and without cell rearrangements (Fig. 5). Additionally, in cases without cell rearrangement, we fitted these moduli with the generalized Maxwell model in Eq. (8) while setting *N*_Maxwell_ = 2.

**Fig. 5.**
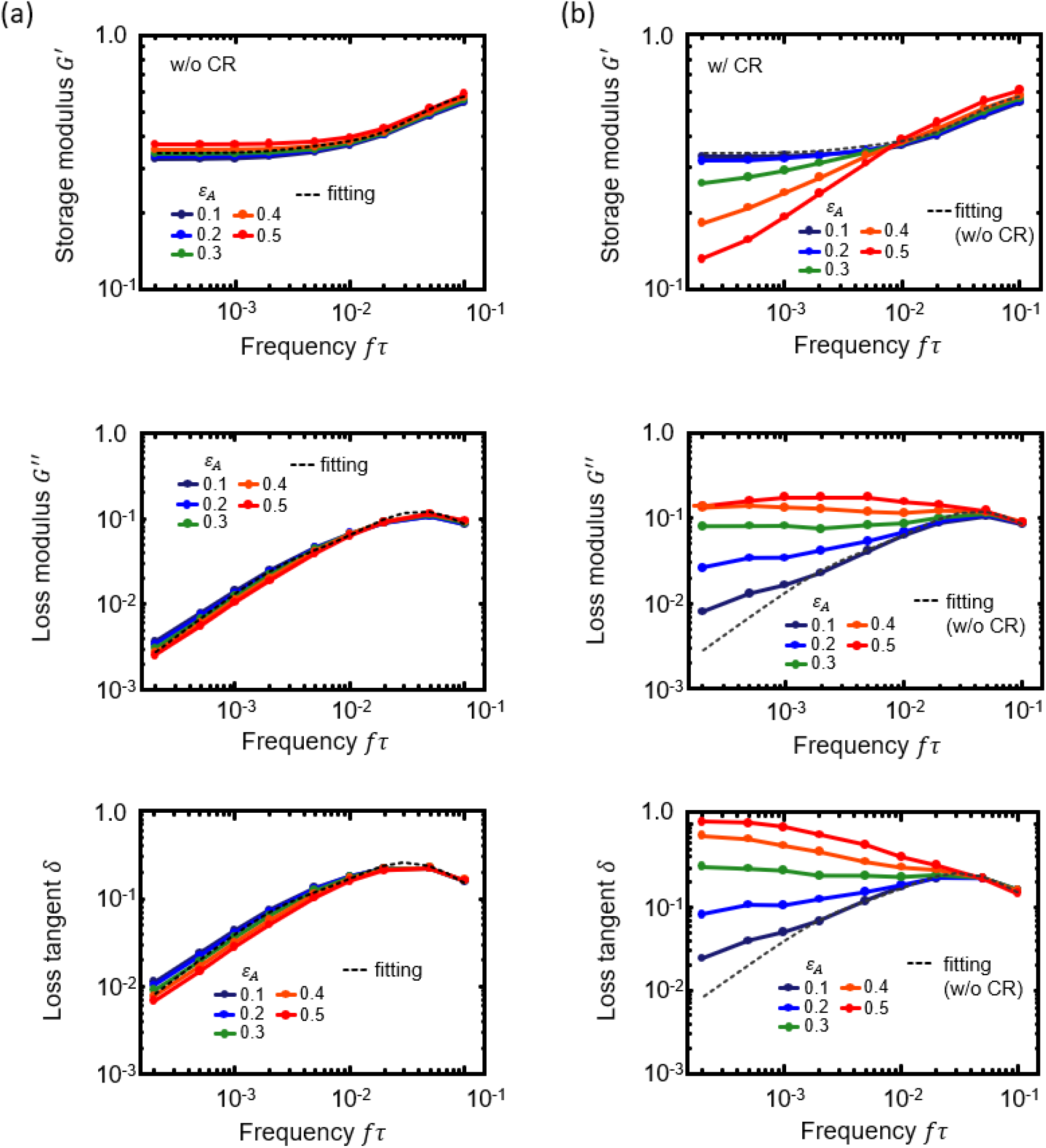
Storage modulus *G′* (top), loss modulus *G ″* (middle), and loss tangent tan *δ*, (bottom) as a function of frequency *fτ* obtained in the dynamic mechanical analysis. Circles connected with straight solid lines indicate different strain amplitudes, *ε*_A_ = 0.1 (black), 0.2 (blue), 0.3 (green), 0.4 (orange), and 0.5 (red). (a) Cases without cell rearrangement (w/o CR). All *ε*_A_ data are fitted to a single curve (dotted line) using the generalized Maxwell model. (b) Cases with cell rearrangement (w/ CR). Additionally, the same dotted lines as in Fig. 5**a** are illustrated to compare data between cases with and without cell rearrangement.

In cases both with and without cell rearrangement (Fig. 5**a**,**b**), *G*′, *G*″, and tan*β* aligned with the Maxwell model at high frequencies (0.02 < *fτ*) but not at low frequencies. In this low-frequency range (*fτ* < 0.02), *G*′ deviated lower than the value predicted by the Maxwell model, while *G*″, and tan*β* deviated higher than the predicted values. These results indicate that the tissue behaves more as a solid during long-term large deformation than that predicted by the Maxwell model. This tendency was more pronounced in the case with cell rearrangement and increased with *ε*_A_.

For example, in the case with *ε*_A_ = 0.5, *G*′, *G*″, and tan*β* highly deviated from the Maxwell model, and their dependence on *f* was significantly altered by cell rearrangement. Particularly, tan*β* decreased linearly with *f*, suggesting that the tissue becomes apparently more fluid with longer timescales (*fτ* < 0.02), which is counterintuitive and differs from the properties of general materials (Fig. 5**b**). However, *G*′ increased linearly with *f* and while *G*″ remains constant with respect to *f*, indicating that the tissue does not become more fluid but becomes less solid at a longer timescale. Thus, the apparent fluidization is caused by the decrease in *G*′ with *ε*_A_, corresponding to strain softening. These findings show that 3D confluent cell tissues exhibit apparent fluidization due to strain softening during long-term large deformation.

A recent study estimated the viscoelastic properties of 3D multicellular tissues using microdroplet magnetic actuation, revealing different behaviors depending on the time scale, i.e., solid-like responses in the short-term and fluid-like responses in the long term [5]. Therefore, our simulation results in the case with cell rearrangement align with this recent study. These findings highlight the utility of the developed method to analyze the dynamic properties of 3D confluent cell tissues.

### 3.4 Multicellular structure and rearrangement induce strain softening and hysteresis of stress-strain relationships

To investigate viscoelastic behavior across different timescales, we calculated the stress *σ* as a function of strain *ε* during dynamic viscoelastic analyses at frequencies *fτ* = 0.001, 0.01, and 0.1 (Fig. 6**a**). For the short timescale (*fτ* = 0.1), both the conditions with and without cell rearrangement exhibited similar responses showing hysteresis, indicated by the area enclosed by the oval trajectory of the *σ*–*ε* relationship (red and black lines in Fig. 6**a-(iii)**). However, as the timescale lengthened (*fτ* = 0.01 and 0.001), hysteresis gradually increased under conditions with cell rearrangement, whereas it decreased and nearly disappeared in the absence of cell rearrangement.

**Fig. 6.**
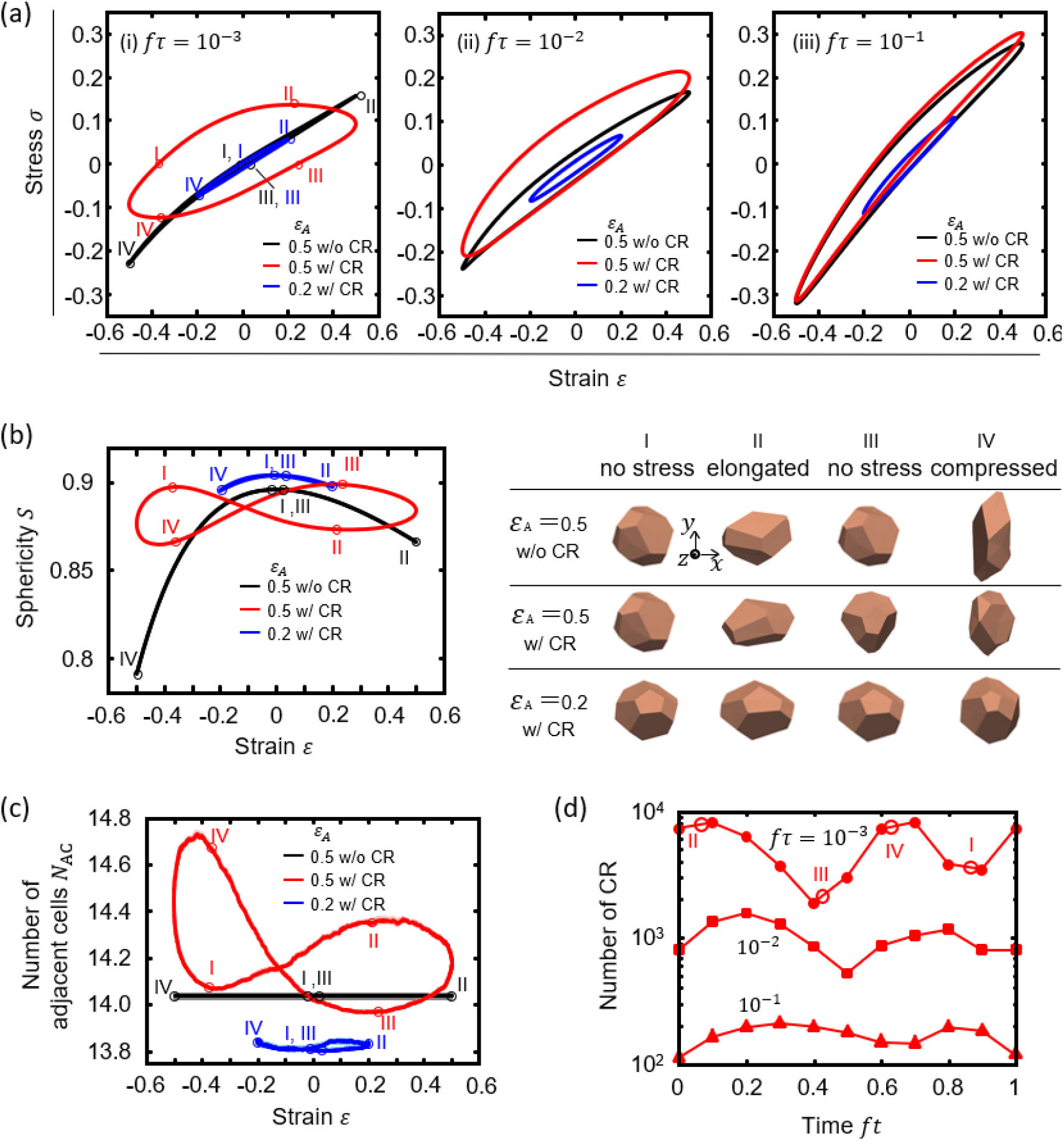
Phase diagrams of (a)-(i) stress, (b) sphericity *S*, and (c) the number of adjacent cells *N*_AC_, as a function of strain *ε*, illustrating hysteresis, which appeared in the dynamic mechanical analysis at low strain frequency (*fτ* = 0.001). Fig. 6**a** also shows phase diagrams of stress *τ* at *fτ* = 0.01 (ii) and (iii) as the middle and high strain frequency cases, respectively. Line colors in Fig. 6**a–c** indicate cases without cell rearrangement (black) and with cell rearrangement (red) at strain amplitude, *εA* =0.5 and with cell rearrangement at *εA* =0.2 (blue). The band widths of light colors indicate the standard deviation. (d) Number of cell rearrangements as a function of time *ft* for strain frequencies *fτ* (circles), 0.01 (squares), and 0.1 (triangles) at strain amplitude *εA* =0.5. The number of cell rearrangements is defined as the total number of cell rearrangements in the entire tissue within the time from *ft* − 0.05 to *ft* + 0.05. For the low-frequency case (*f τ*= 0.001) in Fig. 6**a–d**, the four deformation states are marked as I for *σ=0* in the process from compression to tension, II for the maximum *σ*, III form *σ=0* in the process from tension to compression, and IV for the minimum *σ*. In Fig. 6**b**, the representative cellular shapes are also displayed for the four states I, II, III, and IV.

To investigate the mechanisms producing strain softening and hysteresis during long-term deformation, we focused on the viscoelastic response with *fτ* = 0.001 (Fig. 6**a-(i)**). Additionally, we examined cell-level dynamics by calculating the sphericity *S* and the number of adjacent cells *N*_AC_ as functions of strain *ε* (Fig. 6**b–c**). Each of the *σ*–*ε, σ*–*S*, and *σ*–*N*_AC_ relationships exhibited dynamic transitions with cyclic loops. To discuss the transition processes of these loops, we defined four characteristic states based on the *σ*–*ε* relationship: state I at the transition point from compression to tension with *σ* = 0, state II at the point with the maximum *σ*, state III at the transition point from tension to compression with *σ*= 0, and state IV at the point with the minimum *σ*.

In the case with large deformation and cell rearrangement (*ε*_A_ = 0.5, red line in Fig. 6**a-(i)**), the *σ*–*ε* relationship formed an oval trajectory with a large area, indicating high hysteresis. During the tensile loading from state I to II, *σ* increased with *ε* following a convex upward curve, indicating strain softening as a nonlinear response. From state II to III, *σ* decreased even when *ε*decreased, reaching zero with a positive plastic strain at state III. During this process (0 < *σ*), *S* decreased from state I to II and increased from state II to III (Fig. 6**b**), and *N*_AC_ increased from state I to II and decreased from state II to III (Fig. 6**c**). During compressive loading from state III to IV, *σ* decreased following a convex downward curve, indicating strain softening (Fig. 6**a-(i)**). From state IV to I, *σ* increased even when *ε* decreased, reaching zero with a negative plastic strain at state I. During this process (*σ* < 0), *S* decreased from state III to IV and increased from state IV to I (Fig. 6**b**), and *N*_AC_ increased from state III to IV and decreased from state IV to I (Fig. 6**c**). These behaviors of *S* and *N*_AC_ resulted in a figure-eight shape of the *S*–*ε* and *N*_AC_ –*ε* relationships (Fig. 6**b**,**c**).

To clarify the effects of large strain and cell rearrangement on strain softening and hysteresis, we compared these results with those from conditions with low strain (*ε*_A_ = 0.2, blue lines in Fig. 6**a–c**) and without cell rearrangement (black lines in Fig. 6**a–c**). In the tensile process (state I to II) of the low strain or absence of the cell rearrangement conditions, the increase in *σ* led to a decrease in *S*. In the tension relaxation process (state II to III), the relaxation of *σ* led to an increase in *S* along the same process from I to II. In the compression process (state III to IV), the decrease in *σ* led to a decrease in *S*. In the compression relaxation process (state IV to I), the relaxation of *σ* led to an increase in *S* along the same process from III to IV. Throughout these processes, *N*_AC_ remained constant. These results show that while the relationships among *σ, S*, and *ε* are qualitatively similar regardless of the deformation amount and presence of cell rearrangement, several significant characteristics emerge from large deformation and cell rearrangement. First, in the case with low strain, *N*_AC_ remained almost constant (blue line in Fig. 6**c**), indicating that cell rearrangements require large deformation. Second, in either the case without cell rearrangement or with small strain (black and blue lines in Fig. 6**a–(i)**), the areas enclosed by *σ*–*ε* curves were very small, indicating low hysteresis, showing that hysteresis is primarily caused by cell rearrangement. Third, in the case without cell rearrangement (black line in Fig. 6**a-(i)**), the *σ*–*ε* relationship forms an almost linear line with slight curvature, suggesting that strain softening is not only caused by cell rearrangement but also inherent in the multicellular structure of 3D confluent cell tissues.

The hysteresis observed in the case with large deformation and cell rearrangement (*ε*_A_ = 0.5, red line in Fig. 6**a-(i)**) can be explained as follows. During the tensile loading from state I to II, individual cells recovered their shapes by their resilience (as discussed in Sec. 3.2) to decrease tensile stress (Fig. 6**b**), and this cell shape recovery caused cell rearrangement to enhance this recovery (Fig. 6**c**). Importantly, this cell rearrangement provided strain change to reduce tensile stress, softening the tissue and resulting in a positive plastic strain when reached zero at state III after unloading tensile stress. This process repeated similarly during the compressive loading and unloading from state III to I. Thus, we demonstrated that cell rearrangement, caused by cell shape recovery, results in hysteresis of 3D confluent cell tissues.

The hysteresis loop was affected by strain frequency *f* (Fig. 6**a**) as follows. *σ* increased with *f*, indicating increasing elastic force as illustrated by *G*′increasing with *f* (Fig. 5). In the case without cell rearrangement, the area enclosed by the *σ*–*ε* curve relative to its perimeter increased with increasing *f* because of the viscosity generated by a large strain rate, as illustrated by the increase in *G*″ with *f* (Fig. 5**a**). However, the relative area enclosed by the *σ*–*ε* curve decreased with increasing *f* in cases with cell rearrangement, where the viscosity *G*″ generated by cell rearrangement decreased (Fig. 5**b**). To clarify this point, we calculated the number of cellular rearrangements as a function of regularized time *ft* across different frequencies *f* (Fig. 6**d**). The number of cell rearrangements increased drastically with rising frequency. Notably, at a low frequency (*fτ* = 0.001), the number of cell rearrangements peaked before the strain reached the maximum value, causing the absolute value of *σ* to reach its maximum. As a result, the hysteresis loops differed between the cases with and without cell rearrangement at low *f* (Fig. 6**a**). This finding explains how viscoelastic moduli *G*′, *G*″, and tan*δ*, depend on cell rearrangement at low *f*.

To clarify the origin of strain softening, we calculated the Fourier transform of stress *σ* (Fig. 7). The imaginary-part and real-part components 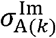 and 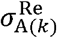 represent in-phase and antiphase components of *σ* as the stress amplitudes, expressing elastic and viscous properties, respectively (Fig. 7**a, b**). Both the stress magnitudes 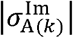 and 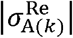 increased with strain *ε*. Moreover, 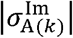 was higher in the case without cell rearrangement than that in the case with cell rearrangement (Fig. 7**a**), whereas 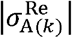 was higher in the case with cell rearrangement than that without rearrangement (Fig. 7**b**). This result indicates that cell rearrangement decreases the elastic response and increases the viscous response, corresponding to the long-term behaviors of *G*′, ″ and tan*δ* (Fig. 5). Additionally, both 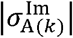 and 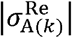 have finite values at *k* < 5, indicating that strain softening can occur irrespective of cell rearrangements but is inherent in multicellular structures.

**Fig. 7.**
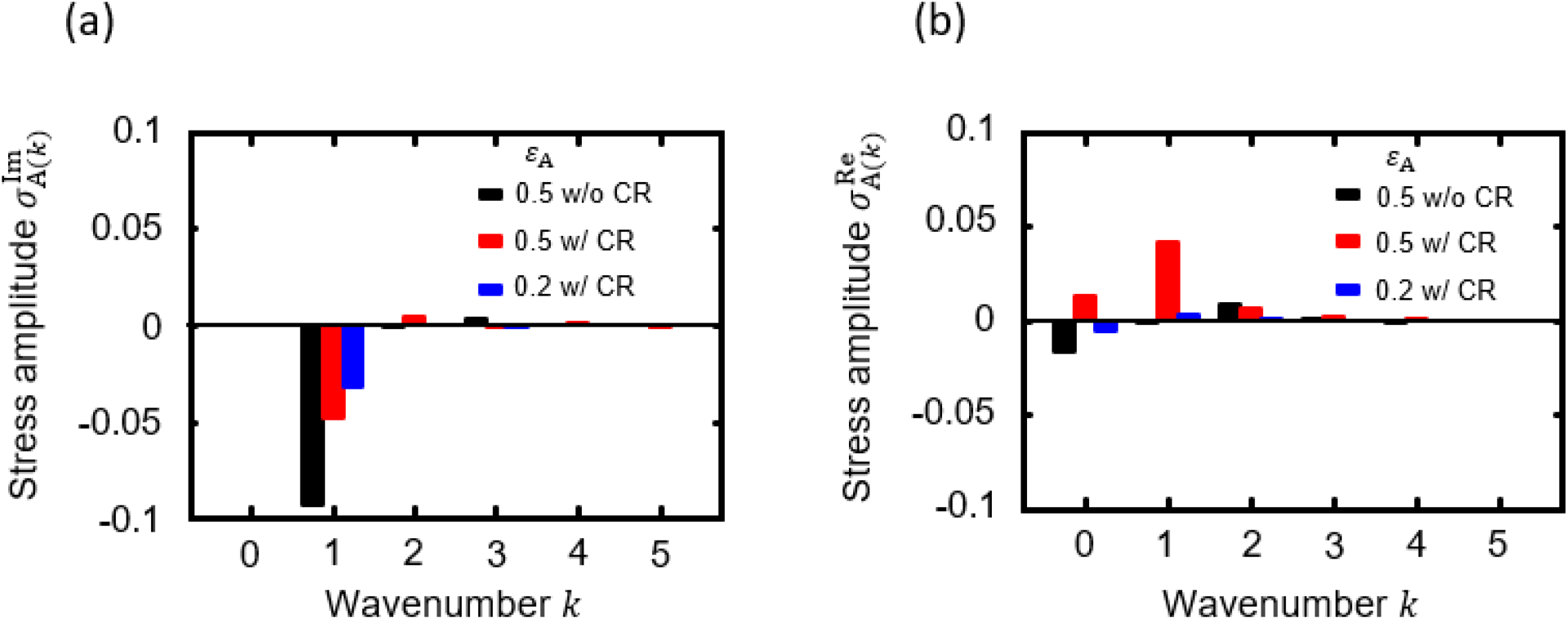
Fourier transform of stress *σ* in the dynamic mechanical analysis in cases without cell rearrangement (w/o CR, black) and with cell rearrangement (w/ CR, red) at strain amplitude *ε* _A_ = 0.5 and with cell rearrangement at *ε*_A_ = 0.2 (w/CR, blue) at strain frequency. *fτ =* 0.001. The frequency components are shown as the stress amplitudes in (a) the imaginary-part (in-phase components to the applied strain *ε*) and as (b) the real part (antiphase components) as a function of wavenumber *k* with respect to *f*.

### 3.5 Ramberg–Osgood model characterization of the long-term properties of 3D confluent cell tissues

To step up from the cell to the organ and body scales, a continuum description of tissue properties is needed. Although the mechanical properties of 3D confluent cell tissues agreed well with the generalized Maxwell model representing the viscoelasticity for small or short-term deformations, they deviated significantly in the case of long-term large deformation (Sec. 3.1–3.3). However, the tissue exhibited elastoplastic deformation over long timescales (Sec. 3.4). To describe the long-term responses of 3D confluent cell tissues as a continuum, we focused on the strain-softening behavior from state I to II obtained in the dynamic viscoelastic analysis in a low-frequency case (Fig. 6**a-(i)**) and introduced the Ramberg–Osgood model [48].

The stress-strain relationships in the deformation process from state I to II at *fτ* = 0.001 for arious strain amplitudes *ε*_A_ (Fig. 6**a-(i)**) were fitted to the Ramberg–Osgood model. The fitted equation is described as

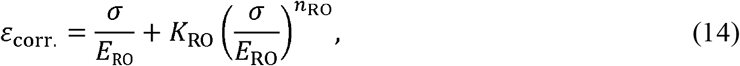

where

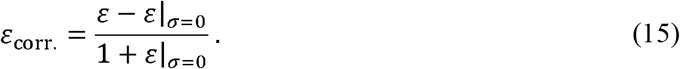

In Eqs. (14) and (15), *ε*_corr_ is the strain corrected to have its reference shape in state I where *σ*, and *ε*|_*σ =* 0_ in Eq. (15) is the strain in state I.

The Ramberg–Osgood model agreed well with the results of the dynamic viscoelastic analyses across a wide strain amplitude *ε*_A_ range in both cases with and without cell rearrangement (Fig. 8**a**,**b**). Specifically, this model accurately represented the strain softening in the case with cell rearrangement (Fig. 8**b**). By fitting the model to the results, we obtained the material parameters *E*_RO_, *K*_RO_ and *n*_RO_ (Fig. 8**c–e**). In the case without cell rearrangement, the strain-softening coefficient *K*_RO_ was nearly zero (Fig. 8**d**), indicating a linear elastic response at the continuum level. In contrast, in the case with cell rearrangement, *K*_RO_ and strain-softening exponent *n*_RO_ were large (Fig. 8**d**,**e**), indicating a strain-softening response at the continuum level. Young modulus *E*_RO_ was not affected by cell rearrangement (Fig. 8**c**), indicating elasticity at small deformations independent from the cell rearrangement and corresponding to the result that large deformation is necessary for cell rearrangement. Additionally, yield strain *ε*_Y_ was calculated using the Ramberg–Osgood equation by assuming 0.2% strain for the yield strength (the inset of Fig. 8**a**). *ε*_Y_ was smaller in the case with cell rearrangement than in the case without cell rearrangement (Fig. 8**f**), indicating a plastic response even at a small deformation regime on the order of 10^−2^ in the strain value. Thus, we demonstrated that 3D confluent cell tissues exhibit a typical elastoplastic response over long timescales, and this elastoplastic property can be expressed by the Ramberg–Osgood model at the continuum level.

**Fig. 8.**
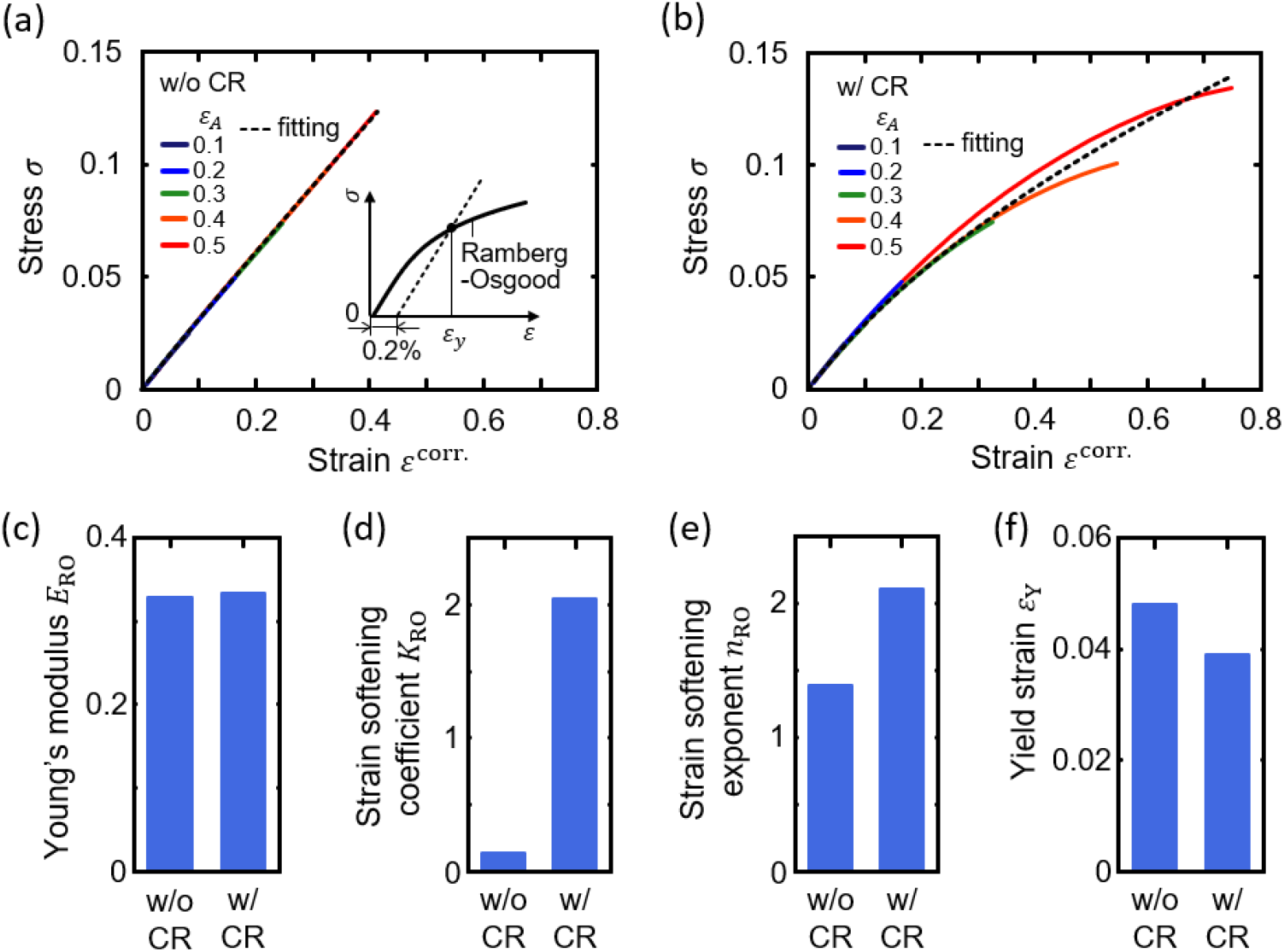
Elastoplasticity in the dynamic mechanical analysis (*f τ* = 0.001) identified using the Ramberg–Osgood equation. (a) In the case without cell rearrangement (w/o CR), stress *σ* between the deformation states I and II (Fig. 6**a–(i)**) is plotted as a function of corrected strain *ε*_corr._ (Eqs. (14) and (15)) for different strain amplitudes *ε*_A_ (solid lines). The stress-strain relationships of all *ε* values were fitted by a single curve with the Ramberg–Osgood equation (dotted lines). The inset diagram illustrates how the yield strain *ε*_Y_ is defined from the Ramberg–Osgood equation by assuming 0.2% strain for the yield strength. (b) The same as in Fig. 8**a** but in the case with cell the rearrangement (w/ CR). (c) Young’s modulus *E*_RO_, the strain-softening (d) coefficient *K*_RO_, and (e) exponent *n*_RO_ involved in the Ramberg–Osgood equation. (f) Yield strain *ε*_Y_ assuming 0.2% strain for the yield strength.

## 4. Discussion

In this study, we developed a new method using a 3D vertex model that enables analysis of the dynamic properties of 3D confluent cell tissues, including long-term large deformations. This method successfully identified characteristic properties arising from multicellular structures and their rearrangements, such as strain softening and hysteresis during long-term large deformations. It also identified that cell rearrangement plays a crucial role in homogenizing the shapes of individual cells. Our method enables a quantitative assessment of the mechanical properties of these tissues at the resolution of individual cell deformations, providing critical insights into tissue mechanics and offering guidance for the development of biomaterials.

During embryogenesis, tissues undergo various deformations due to the activities of their constituent cells and the influence of surrounding tissues [52–54]. Successful morphogenesis requires that tissues retain necessary deformations while resisting detrimental ones. When applying our newly developed method to 3D tissues composed of confluent homogenous cells, we observed that these tissues exhibited viscoelastic and plastic responses (Fig. 2). Specifically, their behavior transitioned from a solid-like elastic response during short-term deformations to a more fluid-like viscoelastic response during long-term deformations (Fig. 6). This transition was significantly more pronounced than the transition seen in the Maxwell model and plays a critical role in balancing the retention and resistance of deformations. This finding is consistent with previous studies that reported elastic responses in tissues, enabling them to resist external forces [55–58], and tissue fluidization, which facilitates structural remodeling [8–10]. Additionally, we found that this transition arises from cell rearrangements, highlighting their significance in achieving morphogenesis. Moreover, the solid-to-fluid transition allows tissues to adapt to deformations, supplemented by other mechanisms such as the irreversibility of epithelial folding induced by an elastoplastic transition, which we previously reported as primarily caused by actin polymerization, rather than cell rearrangement [11]. Such molecular behaviors, including actomyosin contraction and cadherin adhesion, could be integrated with cell rearrangements to endow tissues with more complex properties. These molecular behaviors can be incorporated into the developed method through the energy function of *U*, representing subcellular mechanical behaviors. Key molecular behaviors vary across different tissues and phenomena, and their effects on properties could be better understood and predicted by refining our developed method.

We revealed that multicellular structures and rearrangements provide characteristic properties of 3D confluent cell tissues, especially strain softening and hysteresis during long-term large deformations (Fig. 6). Interestingly, strain softening originates from multicellular structures, while hysteresis is caused by cell rearrangements. Even without cell rearrangement, individual cell-cell boundaries can be flexibly deformed due to the foam-like structure of confluent cell tissues. This cell-cell boundary deformability allows neighboring cells to slide past each other, shifting the tissue response from linear elastic to exhibiting strain softening (Fig. 7). However, in the absence of cell rearrangement, the topological connections among cells are maintained, preventing the entire tissue structure from being recovered solely by individual cell shape recovery. In contrast, with cell rearrangement, the topological connections among cells can be reconfigured, allowing the entire tissue structure to recover through the recovery of individual cell shapes. This process facilitates strain softening and induces hysteresis. Thus, while strain softening does not originate from cell rearrangement, cell rearrangement enhances and amplifies it and transforms the response from elastic to plastic, leading to hysteresis. Importantly, this strain softening is advantageous for multicellular tissues to undergo large deformations during morphogenesis, given the limitations of cellular force generation. Moreover, individual cells constantly generate relatively short-term forces through behaviors such as division, apoptosis, and contraction, which can introduce noise [22,59]. The hysteresis observed only in long-term deformation may play a key role in retaining appropriate deformations, filtering out noise from short-term individual cell forces. These findings highlight the impact of the characteristics emerging from multicellular structures and the difficulty of extrapolating these properties from individual cell measurements. This result indicates that designing mechanical properties of multicellular tissues requires considering the dynamic interactions among cells.

A pioneering study using a 2D vertex model demonstrated linear viscoelastic responses under small strains (*ε* ≤ 0.1) with few cell rearrangements and typical viscoelastic behaviors [60]. Our findings, which are consistent with this previous report, showed small differences between conditions with and without cell rearrangements at low strains (e.g., *ε* ≤ 0.1). In contrast, under large strains (e.g., *ε* ≤ 0.5), we observed significant differences, including strain softening and hysteresis during long-term large deformation, which have not been previously reported. Although it remains unclear whether these nonlinear responses arise from three-dimensionality, there are distinct topological differences in cell movement between 2D and 3D systems. In 3D environments, where cells can deform and move freely in all directions, cells can more easily alter their configurations than with constrained movement within a 2D plane. Further investigation is needed to determine how spatial dimensionality affects rheological properties.

To understand the complex behaviors of tissues comprising numerous cells, particularly during late-stage organogenesis and mature organ functioning, it is essential to examine their long-term behaviors using a continuum description of tissue properties. We revealed that tissues exhibit elastoplastic responses during these long-term deformations, which is well represented by the Ramberg–Osgood model (Fig. 8). This model allows us to coarsely define tissue behaviors, enabling the calculation of long-term, large-scale tissue deformations. Moreover, our simulations provided quantitative parameters for the Ramberg–Osgood model (Fig. 8), illustrating that our method allows us to define continuum models based on specific cellular behaviors, such as actomyosin contraction and cadherin adhesion. However, our method requires further refinement of several points. For example, because it relies on a 3D vertex model that simplifies each cell’s shape into a polyhedron, this simplification may lead to quantitative discrepancies in predicted tissue properties compared with the actual properties, similar to differences between 3D vertex and Voronoi models [61]. These discrepancies could be reduced by adopting more detailed descriptions of cell shapes, similar to those in cellular membrane models [51,62–64]. Given the large number of cells in macroscopic tissues and organs, simulating these tissues with single-cell resolution poses significant challenges. Therefore, multiscale analyses from molecular to organ behaviors are feasible by combining our method, which determines local continuum properties based on molecular behaviors, with broader methods such as finite element methods that model organ behaviors as a continuum [65–68]. By integrating these developed methods with continuum models, we can bridge the gap in understanding from cellular behaviors to organ-level functions.

The mechanical properties of 3D tissues are often overlooked in current biomaterial designs. The methods developed in this study enhance biomaterial engineering by providing a framework for designing tissues with specific mechanical properties. While initially applied to a simple system with homogenous cell properties, the methods can be adapted to more complex systems typical of biological contexts. For example, biased tension at cell-cell boundaries, resulting from actomyosin contractility and cadherin adhesion, can be achieved by modifying the energy function. This approach is also applicable to systems with heterogeneous cell properties and states, such as varying cell-cell boundary tensions and volume elasticity. Extending the model will enable analysis of how these cellular behaviors affect tissue properties, contributing to the design of engineered biomaterials.

## 5. Conclusion

This study proposed a new method to analyze the mechanical properties of 3D multicellular tissues using a 3D vertex model framework. This method offers significant advantages by enabling the assessment of tissue properties with single-cell resolution in response to a wide range of deformation magnitudes and across various time scales. Through a series of analyses, we successfully characterized the viscoelastic and elastoplastic properties of 3D tissues composed of confluent homogeneous cells. Notably, dynamic viscoelastic analyses revealed strain softening and hysteresis under large deformations, arising from cellular structures and rearrangements, respectively. Our method provides a versatile tool to characterize the dynamic properties of 3D multicellular tissues and could serve as a powerful tool for understanding and designing biomaterials.

## Acknowledgments

We would like to thank Tomotaka Fukatsu and Jumpei Kanno, former members of the Tsubota laboratory at Chiba University, for supporting analyses and discussions. This work was supported by the Japan Science and Technology Agency (JST), CREST [Grant No. JPMJCR1921, JPMJCR24B2]; the Japan Agency for Medical Research and Development [AMED; Grant No. 23bm0704065h0003]; the Japan Society for the Promotion of Science (JSPS), KAKENHI [Grant No. 24H01937, 24H01398, 22H05170, 21KK0134, 21H01209, 23K26032, 23H01337]; the World Premier International Research Center Initiative, MEXT, Japan.

## Author Contribution Statement

KT and SO conceived the project and wrote the manuscript; SH, KT, and SO developed software and performed the analyses; TH provided crucial ideas. All authors contributed to the final manuscript.

## Data Availability Statement

The datasets analyzed during the current study are partially available from the corresponding author upon reasonable request.

## Notes

### Competing Interest Statement

The authors have declared no competing interest.

